# Single-cell transcriptome profiling of the *Ciona* larval brain

**DOI:** 10.1101/319327

**Authors:** Sarthak Sharma, Wei Wang, Alberto Stolfi

## Abstract

The tadpole-type larva of *Ciona* has emerged as an intriguing model system for the study of neurodevelopment. The *Ciona intestinalis* connectome has been recently mapped, revealing the smallest central nervous system (CNS) known in any chordate, with only 177 neurons. This minimal CNS is highly reminiscent of larger CNS of vertebrates, sharing many conserved developmental processes, anatomical compartments, neuron subtypes, and even specific neural circuits. Thus, the *Ciona* tadpole offers a unique opportunity to understand the development and wiring of a chordate CNS at single-cell resolution. Here we report the use of single-cell RNAseq to profile the transcriptomes of single cells isolated by fluorescence-activated cell sorting (FACS) from the whole brain of *Ciona robusta* (formerly *intestinalis Type A)* larvae. We have also compared these profiles to bulk RNAseq data from specific subsets of brain cells isolated by FACS using cell type-specific reporter plasmid expression. Taken together, these datasets have begun to reveal the compartment- and cell-specific gene expression patterns that define the organization of the *Ciona* larval brain.

## Introduction

Determining the genetic and cellular bases of an animal’s behavior requires identifying and characterizing all the neurons that comprise its nervous system and understanding how they connect to one another to function in specific neural circuits. The *Ciona* larval nervous system has emerged as an intriguing model in which to study these processes. *Ciona* embryos have long been valued as a developmental model, buoyed by their numerous experimental advantages like small size, low cell number, stereotyped cell lineages, rapid development, compact genome, and their amenability to electroporation with plasmid DNA (Zeller, 2018). *Ciona* also shows largely untapped potential as a model organism for neuroscience. The complete connectome of the 177 central nervous system (CNS) and 54 peripheral nervous system (PNS) neurons of the *Ciona* larva has been recently described in thorough detail by serial electron microscopy (Ryan et al., 2016, 2017, 2018). This is only the 2^nd^ complete connectome mapped, after the nematode *Caenorhabditis elegans*, and one of the smallest nervous system described in any animal (231 neurons in *Ciona* vs. 301 neurons in *C*. elegans)(White et al., 1986). The fact that *Ciona* belongs to the tunicates, the sister group to the vertebrates, makes this minimal nervous system a unique model in which to study chordate-specific principles of neurobiology and neurodevelopment (Nishino, 2018).

Key to understanding the development of multicellular embryos and organs like the brain is the ability to assay gene expression in specific cells or cell types. Transcriptome profiling by DNA microarrays or high-throughput sequencing has proved to be a very powerful tool for such assays, especially given the invariant cell identities and lineages of the *Ciona* embryo, and the ease with which cells can be isolated, for instance by fluorescence-activated cell sorting (FACS). Profiling defined cell populations experimentally isolated from dissociated *Ciona* embryos has been extensively (Christiaen et al., 2008; José-Edwards et al., 2011; Racioppi et al., 2014; Razy-Krajka et al., 2014; Reeves et al., 2017; Wagner et al., 2014; Woznica et al., 2012) and has been instrumental in gaining a whole-genome understanding of gene regulation during development, including in the nervous system (Hamada et al., 2011). However, this approach only detects average gene expression across the entire cell population, as a single cDNA library is prepared from RNA extracted from pooled cells. This may result in missing significant variation between individual cells within the population, in which unidentified subsets of cells may have very distinct transcriptional profiles. This method is also confounded by contaminating cells-cells that are sorted together with the target population but are transcriptionally distinct from it. Transcriptome profiling has also been performed on individual, dissociated blastomeres from early *Ciona* embryos (Ilsley et al., 2017; Matsuoka et al., 2013; Treen et al., 2018), which has allowed for a cell-by-cell, stage-by-stage, whole-genome view of early development. However, this method is not possible in later development, where cells are too small to be manually isolated or identified prior to RNA extraction and cDNA library preparation.

Recent developments in single-cell RNAseq technology (scRNAseq) have further enhanced the tractability of *Ciona* for developmental studies. scRNAseq is unique in that it allows for *in silico* isolation, identification, and characterization of distinct cell populations based on massively parallel sequencing of transcriptome libraries prepared from thousands of individual cells (Moris et al., 2016; Tanay and Regev, 2017; Trapnell, 2015). Because scRNAseq analysis algorithms allow for *a posteriori* identification of each cell within the population, one can process heterogeneous cell populations, whether this heterogeneity is intentional or not. This allows for discovery of previously unknown cell types or resolving different temporal stages of a single cell type’s developmental trajectory. Indeed, scRNAseq was recently used to reconstruct single-cell developmental trajectories of entire vertebrate embryos (Briggs et al., 2018; Farrell et al., 2018; Wagner et al., 2018). In *Ciona* specifically, scRNAseq was used to monitor spatio-temporal transcriptional dynamics in the cardiopharyngeal mesoderm (Wang et al., 2017).

Here we present our results on the use of scRNAseq to profile cells isolated by FACS from the brains of hatched *Ciona robusta* (formerly *intestinalis Type A)* larvae. After using the 10X Genomics Chromium system to obtain 2607 high-quality single-cell transcriptomes from our sample, we were able to classify cells into various clusters based on similar gene expression profiles using the scRNAseq analysis R package Seurat (Satija et al., 2015). Although significant contamination from endoderm and epidermis was observed in our dataset, these cells were easily identified and analyzed separately from bonafide brain cells and other neurons. As a result, we have generated single-cell transcriptional signatures and differential gene expression data for subsets of *C. robusta* larval brain, peripheral nervous system, epidermis, and even endoderm and mesenchyme. These data, which we are fully releasing to the tunicate community, will provide a foundation for future studies on the development and function of these various cell types and tissues.

## Results

Larval brain cells were labeled by expression of electroporated *Fascin>tagRFP* plasmid (**Figure 1A**). Cells from dissociated, electroporated larvae were isolated using FACS by selecting tagRFP+ cells and counterselecting unwanted mesenchyme cells labeled by *Twist-related.c>eGFP*. ~10,0000 isolated cells were processed by the 10X Genomics Chromium system for single-cell library preparation and libraries were then sequenced on an Illumina Hiseq 2500 system. After quality control (see **Material and methods** for details), the final dataset comprised of 2607 cells with 13,435 genes detected across them, out of 15,288 total genes annotated in the *C. robusta* genome. Within this subset of cells, we found a total of 1949 highly variable genes, which were used to perform linear dimensional reduction by PCA and clustering as implemented in Seurat. After clustering the cells into 10 clusters as visualized by t-distributed stochastic neighbor embedding (t-SNE)(**Figure 1B**), we surveyed the top differentially-expressed genes that distinguish each cluster (**Supplemental Table 1**). These data were then used to pinpoint the putative identity each cell cluster based on prior knowledge of the expression patterns of certain marker genes, guided by previous studies on gene expression data extracted from the ANISEED database (Brozovic et al., 2017).

**Figure 1.**
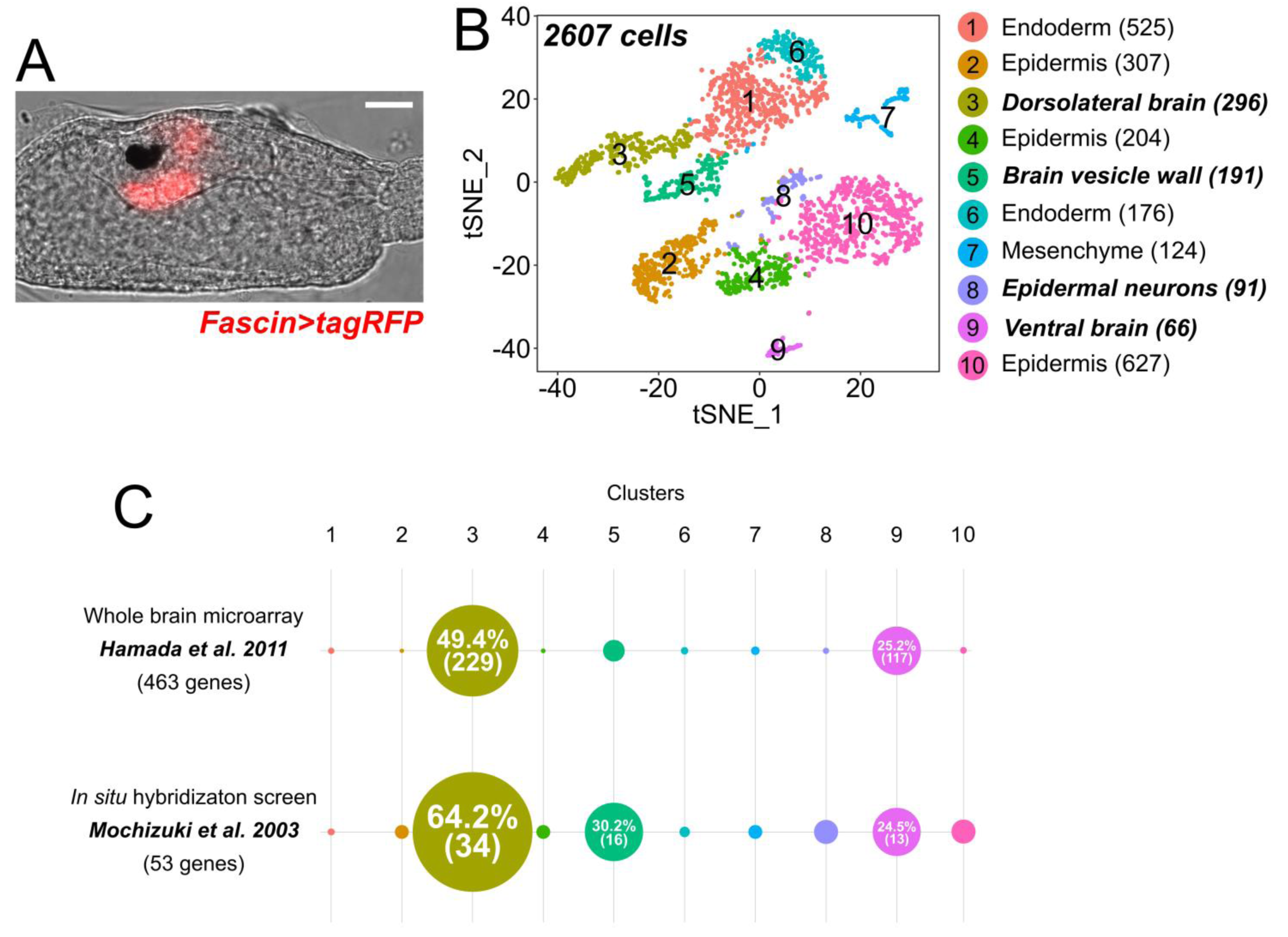
Single-cell RNAseq of sorted *Fascin>tagRFP+* cells. **A)** *Ciona robusta* larva electroporated with *Fascin>tagRFP*, labeling brain cells. Scale bar = 25 μm. **B)** tSNE plot showing all 2607 cells that passed quality control (see **Materials and methods** for details), clustered by Seurat into 10 clusters (1-10). Clusters were assigned putative identities based on gene expression data publicly available on ANISEED. Number of cells in each cluster is indicated in parentheses. Clusters containing mostly neural cells are indicated in ***bold italic*. C)** Comparison of gene expression profiles of Clusters 1-10 to previously published lists of genes enriched in *Ciona* brain (Hamada et al. 2011) and nervous system (Mochizuki et al. 2003). Colored circles indicate the fraction of genes from each previously published study that is also enriched in the corresponding cluster from the present study. Numbers in parentheses indicate total number of genes in each fraction (see **Supplemental Table 2** for details).

Out of these 10 initial clusters, we focused primarily on clusters that clearly represented neural tissue. Based on marker gene expression (explained in more detail below), these were identified as: dorsolateral brain (Cluster 3), sensory/brain vesicle wall (Cluster 5), epidermal neurons (Cluster 8), and ventral brain (Cluster 9). This was confirmed by comparing the list of top differentially expressed genes in each cluster to genes identified in previous surveys of brain or CNS gene expression (**Supplemental Table 2**)(Hamada et al., 2011; Mochizuki et al., 2003). A previous study based on bulk RNAseq profiling of whole Kaede+ brains isolated by partial dissociation of transgenic larvae had identified 565 genes that are preferentially expressed in the brain (Hamada et al., 2011). Of these genes, we were able to determine the KyotoHoya gene identifiers (KH IDs) for 463, which allowed us to survey their enrichment in our neural clusters (**Figure 1C**). Indeed, the majority were found to be expressed in one or more neural clusters, especially Cluster 3, which is overwhelmingly composed of differentiated neurons. Similarly, a set of neuron-expressed genes identified by *in situ* hybridization also had great overlap with those genes enriched in our clusters (Mochizuki et al., 2003). Together, these comparisons indicate that our scRNAseq analysis was able to correctly identify neural cells and capture their relevant neural-specific gene expression.

Of the three major brain clusters (**Fig 2A**), Cluster 3 cells mostly expressed well studied markers of differentiated neurons and photoreceptors associated with the dorsal and lateral regions of the brain. These markers include, *Synaptotagmin 1* (Mochizuki et al., 2003), *Cel3/4/5* (also known as Etr-1)(Satou et al., 2001), and *Rlbp1* (also known as Cralbp)(Mochizuki et al., 2003)(**Fig 2B**)(**Supplemental Table 1, sheet 3**). These are cells that likely descend from the cells marked by the earliest expression of *Fascin>tagRFP* in the lateral rows of the anterior neural tube (**Supplemental Figure 1A**), and are derived from the A9.14/A9.16 blastomeres of the neural plate (Cole and Meinertzhagen, 2004; Imai et al., 2009; Navarrete and Levine, 2016; Oonuma et al., 2016).

**Figure 2.**
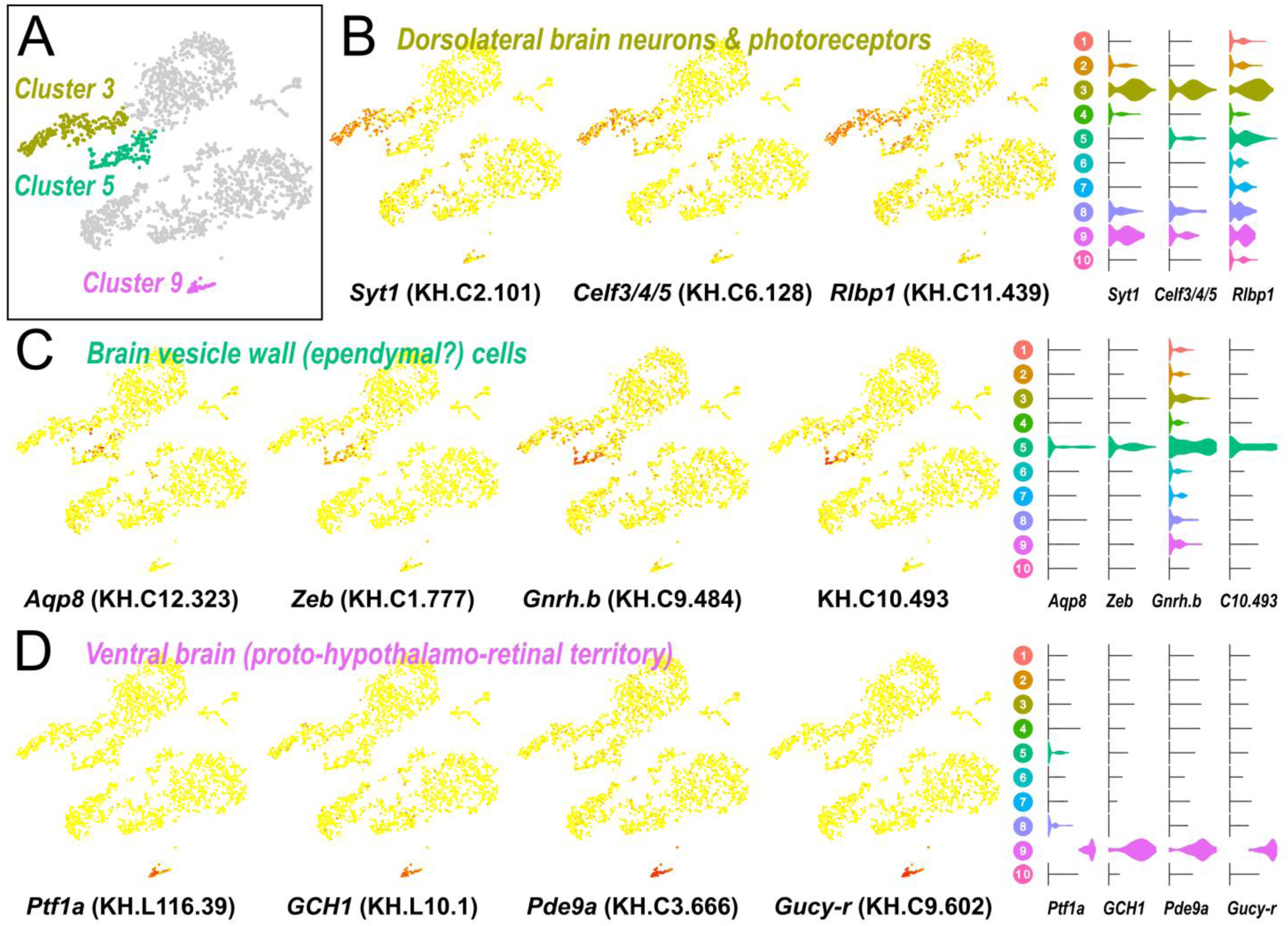
Marker genes reveal cell identities of major brain regions. **A)** Clusters 3, 5, and 9, corresponding to the majority of the *Ciona* larval brain, highlighted in the original tSNE plot (see **Figure 1**). Other clusters and outlier cells are shaded in gray. **B)** Gene expression of neuronal markers *Synaptotagmin 1 (Syt1), Celf3/4/5* (also known as *Etr-1), Rlbp1* (also known as *Cralbp)* plotted on tSNE plots of clustered cells. High expression of neuronal markers in these cells delineates Cluster 3, comprised of neurons and photoreceptors mostly in the dorsolateral regions of the brain. **C)** Cluster 5 cells show expression of brain vesicle wall (ependymal) cell markers *Aquaporin 8 (Aqp8), Zeb, Gonadotropin-releasing hormone.b (Gnrh.b*, previously named *GnRH2)* and gene KH.C10.493. D) Cluster 9 is distinctly marked by expression of *Ptf1a, GTP cyclohydrolase 1 (GCH1) Phosphodiesterase 9a (Pde9a)* and *Guanylate cyclase-related (Gucy-r)*. According to Razy-Krajka et al. (2015), these cells, comprising the ventral-most part of the brain, represent the tunicate homologue of a proto-hypothalamo-retinal region in the olfactorian ancestor that gave rise to both hypothalamus and retinal neurons in vertebrates. In all gene expression maps, red color indicates high expression, yellow indicates low expression. Gene expression violin plots (right) are color coded according to clusters in Figure 1.

Cluster 5 cells clustered closely with dorsolateral brain neurons (Cluster 3) and also expressed *Rlbp1*, but they showed little expression of differentiated neuronal markers (**Supplemental Table 1, sheet 5**). Expression of markers like *Aquaporin 8* (Hamada et al., 2007), *Zeb* (also known as *Zinc Finger (C2H2)-33* or Ci-ZF266)(Imai et al., 2004), and *Gonadotropin-releasing hormone.b* (*Gnrh.b*, previously called GnRH2)(Kusakabe et al., 2003; Kusakabe et al., 2012)(**Fig 2C**), indicated that these might be the so-called ependymal cells lining the sensory or brain vesicle, a specialized fluid-filled cavity important for the function of associated sensory organs such as the ocellus and otolith (Tsuda et al., 2003b). In addition to forming the epithelium that lines the brain vesicle and other parts of the neural tube (Katz, 1983; Mackie and Bone, 1976), these cells also function as neural progenitors that give rise to the adult CNS after metamorphosis (Horie et al., 2011). Expression of *Gnhr.b* and also a known marker of ependymal cells of the tail nerve cord (KH.C10.493)(Hudson et al., 2015; Mochizuki et al., 2003) suggests that this cluster either contains ependymal cells from other parts of the neural tube, captured due to leaky expression of *Fascin>tagRFP*, or a pan-ependymal regulatory program that is shared among similar cells along the entire length of the neural tube.

We also identified a cluster of cells (Cluster 9) that was molecularly very distinct from all other cells. Looking at the top genes associated with these cells, we classified them as cells of the ventral brain, which had been previously characterized by mRNA *in situ* hybridization (Shimozono et al., 2010) and characterized as a “proto-hypothalamo-retinal” territory, hypothesized to be homologous to both the hypothalamus and the retina of vertebrates (Razy-Krajka et al., 2012). These markers include *Ptf1a* and *GTP cyclohydrolase 1* (Razy-Krajka et al., 2012), *Phosphodiesterase 9A* (Shimozono et al., 2010), and *Guanylate cyclase-related* (Hudson et al., 2003)(**Fig 2D**)(**Supplemental Table 1, sheet 9**). These cells arise mostly from a-line lineages (Cole and Meinertzhagen, 2004) and therefore must activate the *Fascin* reporter at a later stage than the A9.14/A9.16-derived cells.

In addition to these three major brain areas, we unexpectedly detected a small cluster of cells (Cluster 8) that was distinguished by a number of cells expressing markers of epidermal sensory neurons and palp neurons (**Supplemental Table 1, sheet 8**).They clustered closely to cells in clusters 2, 4, and 10, which appeared to represent various epidermal cells (**Supplemental Table 1, sheets 2, 4, & 10**). As the name implies, epidermal neurons (as well as palp neurons) are embedded in the epidermis and arise from boundary regions that give rise to both neurons and epidermal cells (Abitua et al., 2015; Gainous et al., 2015; Imai and Meinertzhagen, 2007; Takamura, 1998; Wagner and Levine, 2012). This unintended contamination of epidermal cells and embedded sensory neurons is likely due to leakiness of *Fascin>tagRFP* that was not counter-selected by *Twist-related.c>eGFP*. Indeed, a *Fascin>H2B::GFP* reporter revealed very low-level expression in epidermal cells that may have been picked up by FACS but not by eye (**Supplemental Figure 1B**).

Finally, clusters 1 and 6 were marked by specific expression of known endoderm markers, with Cluster 6 possibly representing more specifically posterior endoderm (**Supplemental Table 1, sheet 1 & 6**), while Cluster 7 expressed markers of mesenchyme (**Supplemental Table 1, sheet 7**), indicating additional unexpected contaminant cells. Although our focus was not on endoderm or mesenchyme, our scRNAseq data provide a glimpse into the transcriptomes of these tissues as well. Thanks to the nature of scRNAseq, these cells were easily excluded *a posteriori* and did not impact our subsequent analyses of neural-specific gene expression.

### Second Stage of Clustering

Since we were particularly interested in neurodevelopment, we focused our second analysis on those clusters that we identified as comprising primarily neural tissue: Clusters 3, 5, 8, and 9, comprising a total of 644 cells. Accordingly, we filtered this subset of cells from our original dataset and repeated PCA and clustering on only these cells. This produced a novel set of 8 clusters by Seurat’s automated cluster identification function. However, upon close manual inspection based on known marker gene expression, we manually identified an additional two clusters, resulting in a final set of 10 clusters, which we called A-J (**Fig 3, Supplemental Table 3**).

**Figure 3.**
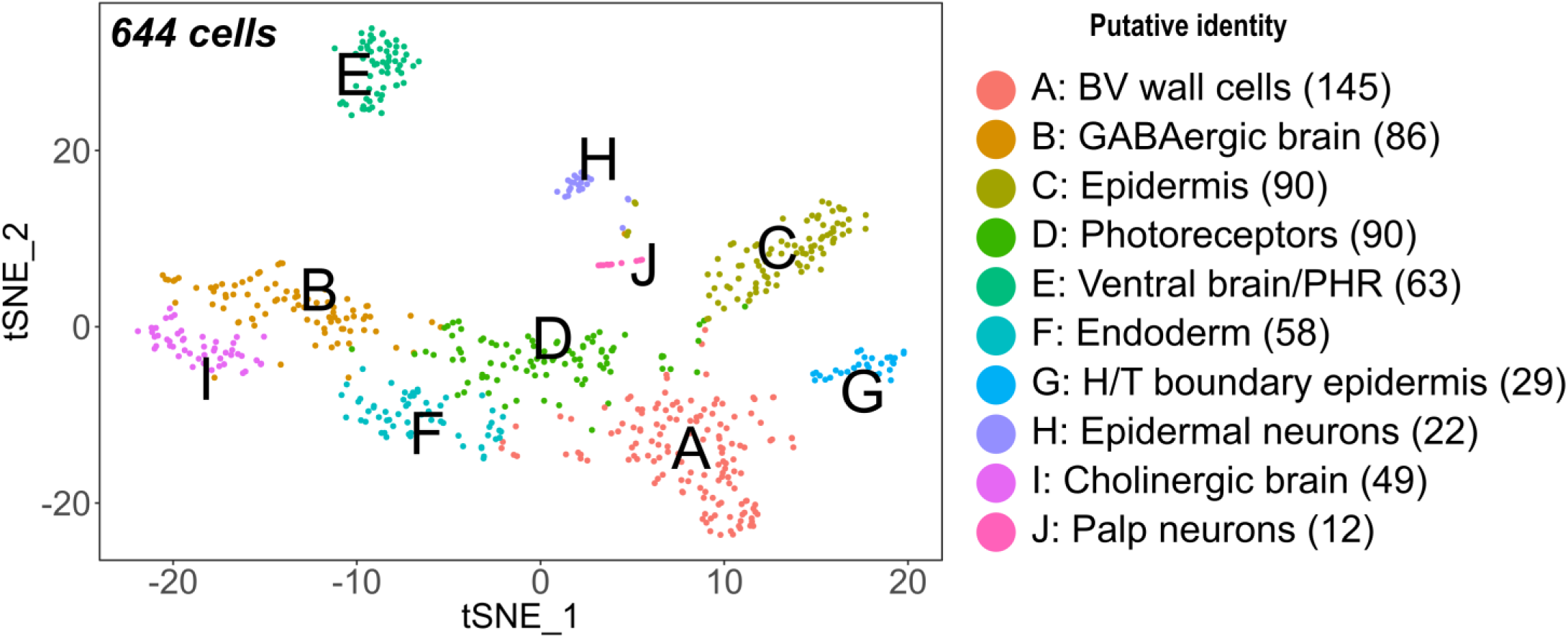
Re-clustering of putative neural cells. tSNE plot showing re-clustering of 644 cells from original clusters 3, 5, 8 and 9, which were deemed to be primarily neural in composition. Cluster I was manually separated from Cluster B, and Cluster J was manually separated from Cluster C. Putative identities are based on known gene expression patterns available on ANISEED (see **Results** for details). BV: brain vesicle, PHR: proto-hypothalamus/retina, H/T: head/tail. Number of cells in each cluster is indicated in parentheses.

### Coronet Cells: Cluster E

In this reclustering, cells of the proto-hypothalamo-retinal region of the ventral brain were grouped in Cluster E (**Supplemental Table 3, sheet 5**). This was largely identical to the previous Cluster 9 and still by far the most clearly delineated, most transcriptionally distinct cells in our analysis. This cluster includes a group of distinctive cells called the coronet cells (Dilly, 1969), which are immunoreactive for dopamine and express *Tyrosine hydroxylase*, which encodes the rate-limiting enzyme for catecholamine synthesis (**Fig 4A,B**)(Moret et al., 2005). To validate this identification, we compared the gene expression profile of Cluster E cells to a coronet cell transcriptome obtained by a more traditional bulk RNAseq profiling method. We isolated *TH>eGFP+* coronet cells from dissociated larvae by FACS. Illumina high-throughput sequencing was performed on total RNA extracted from sorted cells, and analyzed for differential gene expression compared to sorted cells from whole brains labeled by either *Fascin>tagRFP* or *Fascin>Unc-76::eGFP* (**Supplemental Table 4**). We found a high correlation between differential gene expression values from scRNAseq and bulk RNAseq analyses (**Fig 4B, Supplemental Table 5**), suggesting that Cluster E is highly specific for coronet cells and perhaps other closely related cell types of the ventral brain. This comparison validates the use of scRNAseq as an effective means to capture transcriptional signatures of specific cell types in the *Ciona* nervous system.

**Figure 4.**
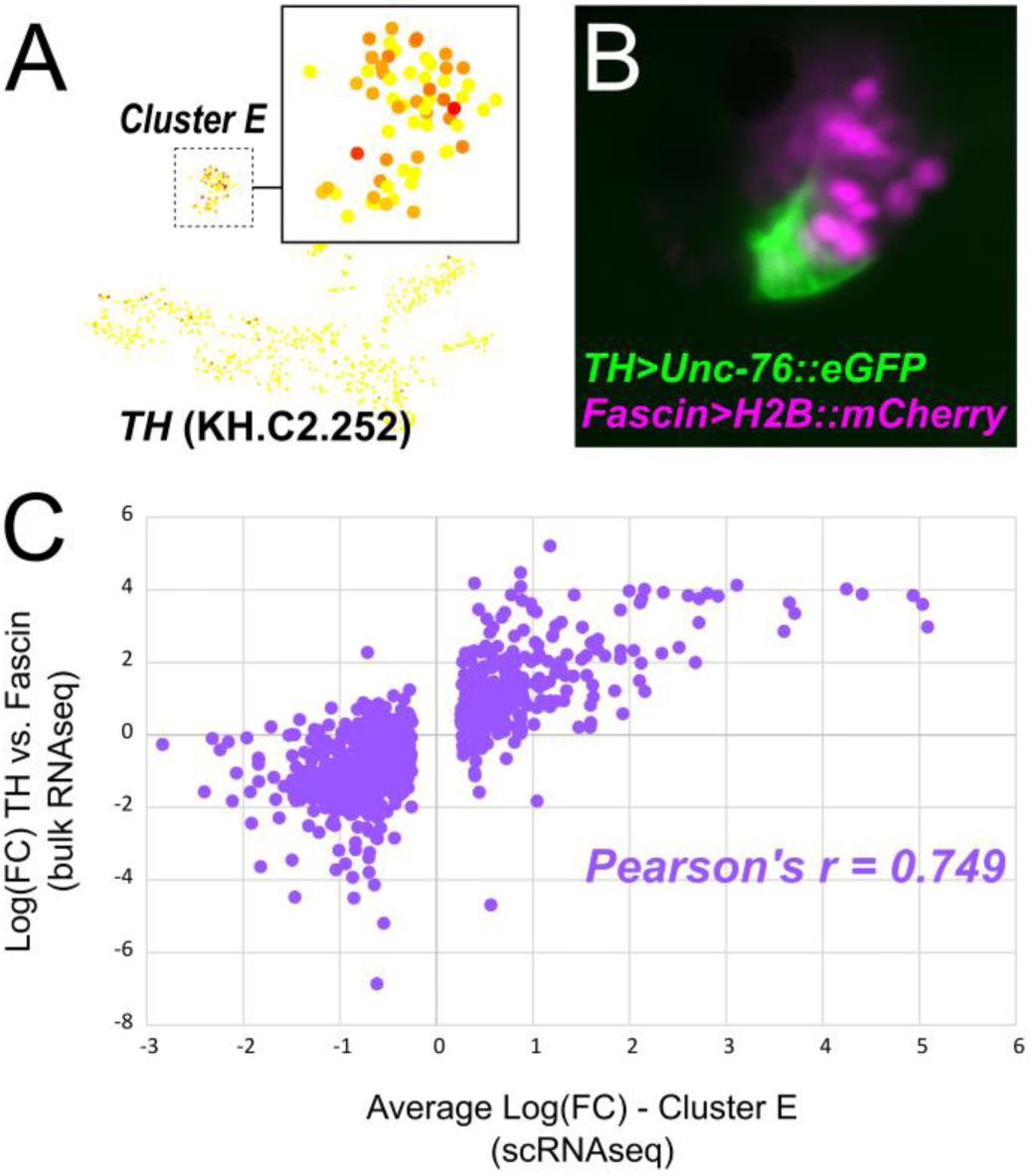
Comparison of gene expression profiling of coronet cells by scRNAseq and bulk RNAseq. **A)** Expression of coronet cell marker *Tyrosine hydroxylase (TH)* mapped onto Cluster E cells on tSNE plot. **B)** Magnified view of brain of *Ciona* larva electroporated with *Fascin>H2B:mCherry* (pink nuclei) and *TH>Unc-76::eGFP* (green), revealing coronet cells of the ventral brain/proto-hypothalamo-retinal region. **C)** Comparison of differential gene expression analyses based on bulk RNAseq (enriched/depleted in sorted TH+ cells relative to whole brain; see **Supplemental Table 4**) and scRNAseq (enriched/depleted in Cluster E, **Supplemental Table 2, sheet 5**), indicating a strong correlation (Pearson’s correlation coefficient, r = 0.749) between the two methods. Only genes that were differentially expressed in Cluster E were analyzed (n = 942 total genes. See **Supplemental Table 5**). “Gap” in scRNAseq values is due to a default average log(FC) cutoff of +/−0.25. For perspective, little correlation was found when comparing values for genes differentially expressed in other neurons, like Cluster B (r = −0.327) or Cluster I (r = −0.271).

The coronet cells are proposed to have evolved from an ancestral neuroendocrine/photoreceptive territory and have been implicated in mediating behavioral responses to light in *Ciona* larvae (Kusakabe, 2017; Razy-Krajka et al., 2012). Furthermore, they are morphologically reminiscent of the coronet cells of the saccus vasculosus in the hypothalamus of fish, which have been shown to have photoreceptive and involved in day length sensing (Nakane et al., 2013). These observations raise the possibility that the coronet cells might be light-sensitive themselves. Expression of the visual cycle gene *Bco1/Rpe65* (KH.C9.224) and the major larval visual Opsin gene, *Opsin1/2.a* (KH.L171.13) is enriched in Cluster 9 cells when compared to all cells in the original dataset (**Supplemental Table 1, sheet 9**), but not when Cluster E cells are compared to the rest of the brain. However, the brain of the *Ciona* larva is relatively rich in photoreceptors. In the connectome, photoreceptors represent the single most numerous cell type in the brain, at a count of 37 out of 143 total cells (excluding ependymal cells)(Ryan et al., 2016). It is likely that this overall overrepresentation of photoreceptors confounds any differential expression of visual cycle genes within specific regions or cell populations within the brain such as the coronet cells. Curiously, our bulk RNAseq data revealed coronet cell-specific expression of a closely related visual *Opsin* paralog, *Opsin1/2.b* (KH.L38.6)(Kusakabe and Tsuda, 2007; Nakashima et al., 2003; Tsuda et al., 2003a), which was filtered from our scRNAseq dataset in quality control. *Opsin1/2.b* expression was only detected in two cells out of entire dataset, and thus did not make the expression cutoff. When we mapped those two *Opsin1/2.b-expressing* cells to our clusters, we found that both belong to Cluster E. Taken together, our data indicate that *Ciona* coronet cells, or a closely related cell type in Cluster E, may indeed be light-sensitive like their vertebrate homologues and the proposed ancestral proto-hypothalamus/retina.

### Peripheral neurons: Clusters H and J

In the original analysis, Cluster 8 appeared to encompass various cells types embedded in the epidermis including epidermal neurons and palps. However, these cells were subsequently resolved into Clusters C, G, H, and J. Of these, Clusters C and G are mostly non-neuronal epidermal cells, based on previous *in situ* hybridization assays (**Supplemental Table 3, sheet 3 and 7**). Cluster G seems to correspond to transcriptionally distinct cells of the head/tail boundary(Kanda et al., 2009), while Cluster C did not, at first glance, appear to comprise any specific compartment of the epidermis. On the other hand, Clusters H and J were clearly neuronal. Cluster H was found to correspond to non-palp epidermal neurons (e.g. RTENs, ATENs, and CENs)(Imai and Meinertzhagen, 2007; Ryan et al., 2018; Takamura, 1998), based on expression of PNS markers such as *Thymosin beta-related* (also called *ϐThymosin-*like)(Pasini et al., 2006)(**Figure 5A,B**) but not *Crystallin beta/gamma*, a marker of palp neurons (**Supplemental Table 3, sheet 8**)(Shimeld et al., 2005). We were also able to resolve a small cluster of 12 palp neurons (Cluster J), which we manually separated from Cluster C based on high *Crystallin beta/gamma* expression (**Figure 5C,D**)(Shimeld et al., 2005). Further analysis of the top differentially expressed genes in this small subset of cells revealed the expression of other previously known palp-specific genes including *Islet* (**Figure 5E,F**)(**Supplemental Table 3, sheet 10**)(Giuliano et al., 1998).

**Figure 5.**
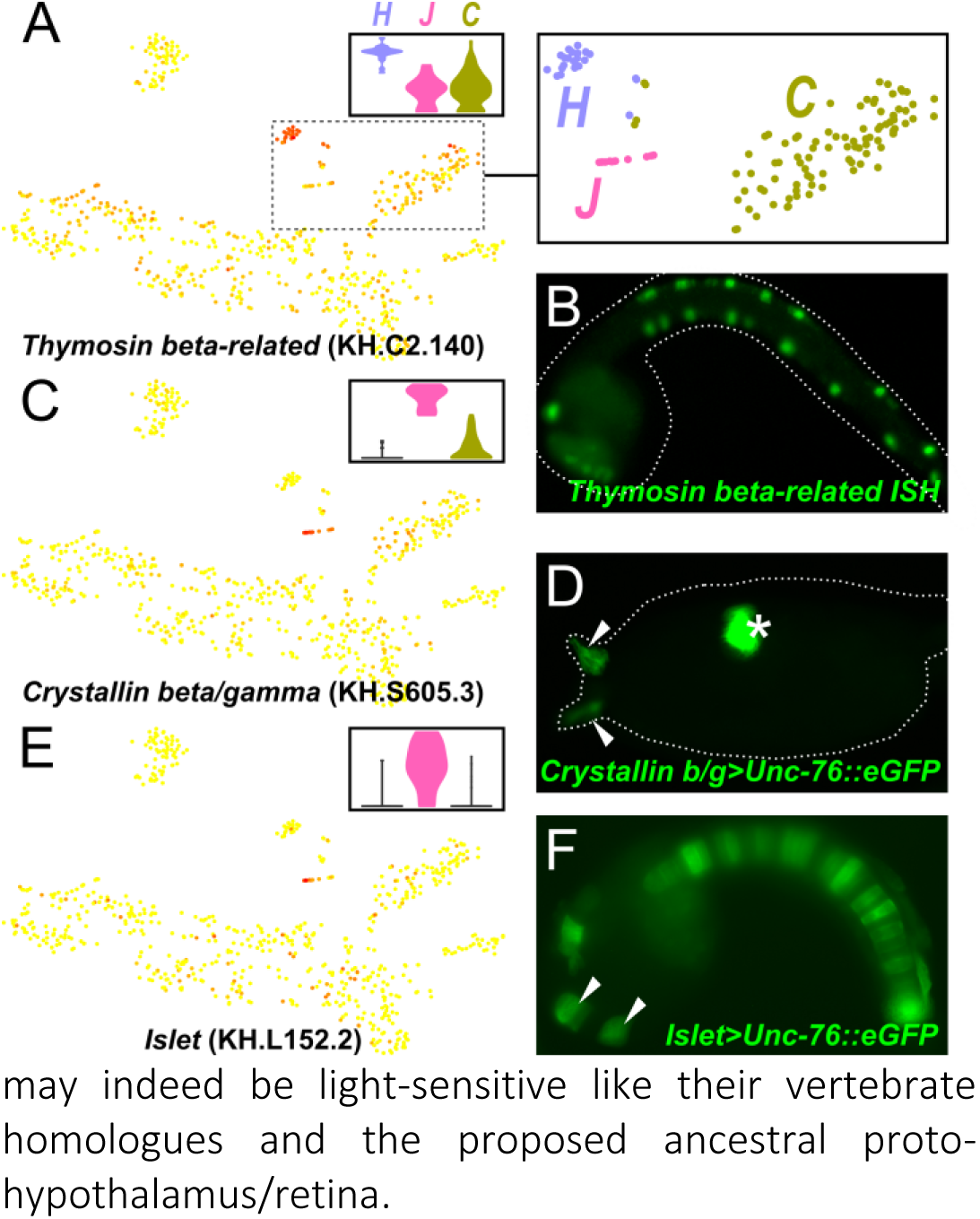
Epidermal and palp neurons. **A)** Gene expression of *Thymosin beta-related* mapped onto the tSNE plot. Dotted outline indicates region magnified in inset at far right, highlighting clusters H, J, and C, excluding certain outlier cells. Top center inset: gene expression violin plots. **B)** *In situ* hybridization of *Thymosin beta-related* in a late tailbud embryo, showing expression in developing epidermal neurons of the head and tail. **C)** Expression of palp neuron marker *Crystallin beta/gamma* marks small 12-cell cluster (Cluster J). Top right inset: gene expression violin plot color coded as in (A). **D)** Larva electroporated with *Crystallin beta/gamma>Unc-76::eGFP* (green), revealing palp neurons (arrowheads). Expression is also strong in the otolith pigment cell (asterisk), which does not express *Fascin* and was therefore not found in our dataset. E) Expression of another known marker of palp neurons, *Islet*, also maps to Cluster J. Top right inset: gene expression violin plot. F) Mid-tailbud embryo electroporated with *Islet[palp+notochord]>Unc-76::eGFP*, labeling developing palp neurons (arrowheads), notochord, and bipolar tail neurons. This particular *Islet* driver, corresponding to the first intron and the proximal promoter region, was identified in Wagner et al. (2014).

### Brain neurons: Clusters B and I

Clusters B and I were initially assigned to a single cluster automatically, but mapping gene expression profiles to these cells clearly identified them as two distinct sub-groups (**Fig 6A** and **Supplemental Table 3, sheets 2 and 9**). Cluster B may correspond largely to GABAergic neurons, based on expression of *Glutamate decarboxylase (GAD)*(**Fig 6B,C**)(Zega et al., 2008), which encodes an essential enzyme for GABA biosynthesis. A subset of these were seen to express *Slc32a1* (also known as *Vesicular GABA transporter*, or *VGAT)*, which encodes the transporter that loads the inhibitory neurotransmitters GABA and glycine into synaptic vesicles (**Fig 6D,E**)(Yoshida et al., 2004).

**Figure 6.**
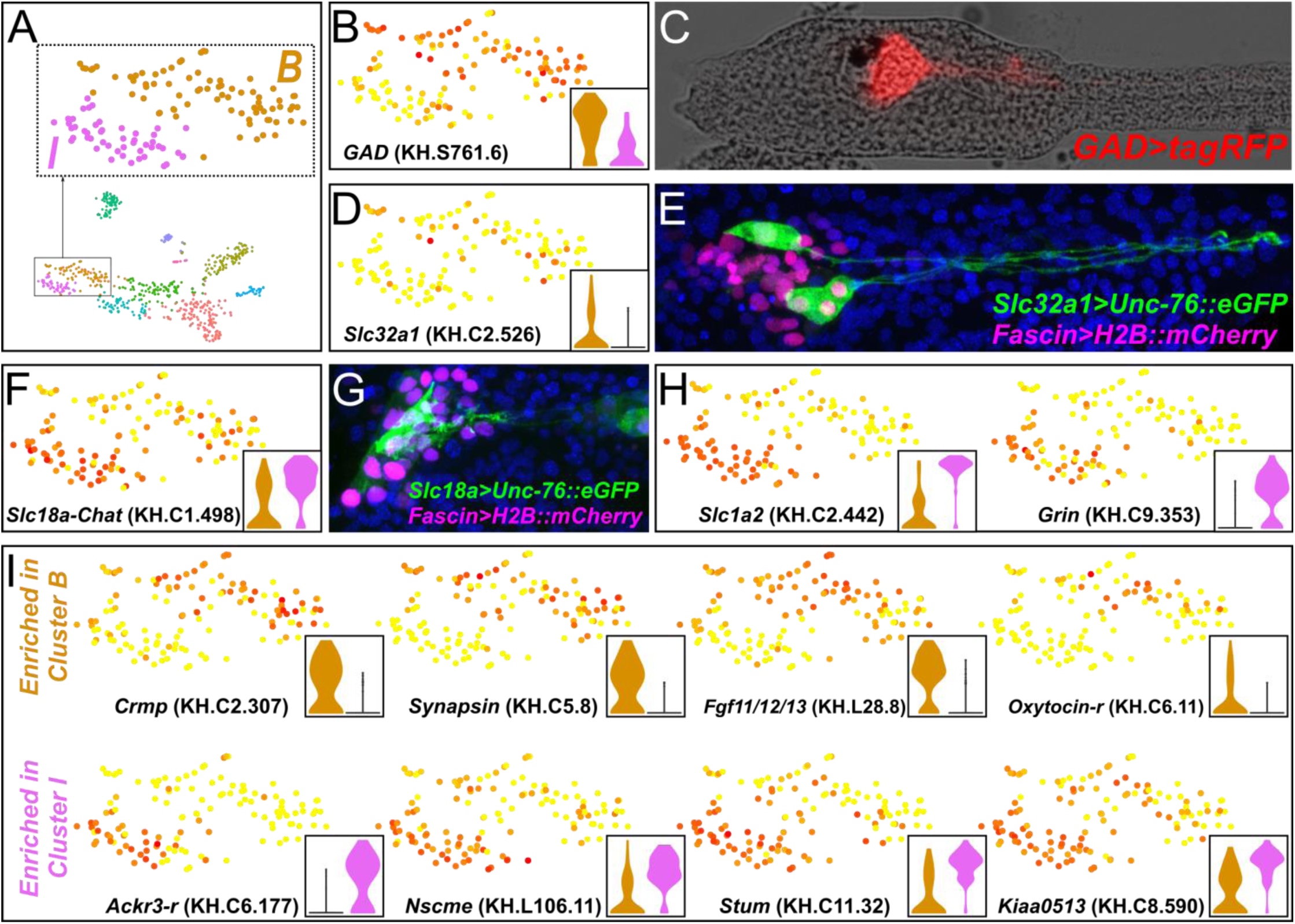
Major subdivision of brain neurons into putative GABAergic and cholinergic neurons. **A)** tSNE plot of neural-specific cells from Figure 3. Inset: enlarged view of Clusters B and I, corresponding to dorsolateral brain neurons, with some outliers omitted. **B)** Expression of *GABA decarboxylase (GAD)* superimposed on clusters B and I, showing enrichment in Cluster B and depletion in Cluster I. Gene expression violin plot in inset, color coded as in (A). **C)** Larva electroporated with *GAD>tagRFP*, revealing *GAD+* neurons of the brain, likely corresponding to Cluster B. **D)** Expression of *Slc32a1* (also known as *Vesicular GABA transporter, VGAT)* in a subset of GAD-expressing neurons. **E)** Larva co-electroporated with *Fascin>H2B::mCherry* (pink nuclei) and *Slc32a1>Unc-76::eGFP* (green), showing *VGAT^+^* brain neurons. All nuclei counterstained with DAPI (blue). **F)** Expression of KH.C1.498 operon, consisting of *Slc18a* (also known as *Vesicular acetylcholine transporter, VAChT)* and *Choline acetyltransferase (Chat)* plotted onto clusters B and I, showing enrichment in Cluster I. **G)** Larva electroporated with *Fascin>H2B::mCherry* (pink nuclei) and *Slc18a>Unc-76::eGFP* (green), showing cholinergic neurons of the brain. All nuclei counterstained with DAPI (blue). **H)** Expression of *Slc1a2* encoding a Glutamate transporter and *Grin*, encoding an NMDA-type ionotropic glutamate receptor, labels primarily Cluster I, suggesting that cholinergic neurons are the principal targets of glutamatergic neurotransmission in the *Ciona* larval brain. **I)** Expression plots of various genes showing enrichment in either Cluster B (top row) or Cluster I (bottom row).

On the other hand, Cluster I appears to correspond to cholinergic neurons, based on expression of *Slc18a* (also known as *Vesicular acetylcholine transporter, VAChT)* and *Choline acetyltransferase (Chat)*, which form a highly conserved operon (KH.C1.498) encoding rate-limiting proteins for cholinergic neurotransmission (**Fig 6F,G**)(Takamura et al., 2002). Interestingly, among the top genes expressed in this group were *Slc1a2* and *Grin* (**Fig 6H**). *Slc1a2* encodes a glutamate transporter protein that clears glutamate from the synaptic cleft, while *Grin* encodes an NMDA-type glutamate receptor. The expression of these genes hints at the intriguing possibility that cholinergic neurons are the major post-synaptic targets of glutamate neurotransmission in the *Ciona* brain. Other genes showed primarily enrichment in either Cluster B or I (**Fig 6I**), suggesting that, among dorsolateral brain neurons the most important distinction may be between excitatory (cholinergic) and inhibitory (GABAergic) cells.

A third class of neurons expressing *Slc17a6/7/8* (also known as *Vesicular glutamate transporter*, or *VGluT)* has been previously reported in the *C. robusta* brain (Horie et al., 2008a). This is the sole *C. robusta* gene encoding the vesicular glutamate transporter protein, which is critical for glutamate loading in the synaptic vesicles of glutamatergic neurons. Its expression was found to be moderately enriched in peripheral neurons (Cluster H, **Supplemental Table 3, sheet 8**), which are known to be glutamatergic, but not in any other clusters. However, we found that the current gene model for *Slc17a6/7/8* (KH.C3.324) is poorly annotated and may have caused undercounting of cells expressing this gene in the brain. In the future, a more accurate gene model for *Slc17a6/7/8* will need to be devised for a reliable characterization of glutamatergic neurons in the *Ciona* brain.

### Photoreceptors: cluster D

Cluster D seemed to contain various cells that were previously clustered with dorsolateral brain neurons in the original Cluster 3. Here, their putative photoreceptor identity is suggested by conspicuous expression of visual cycle genes *Bco1/Rpe65, Opsin1/2.a*, and *Opsin4/Rgr* (KH.L57.28, also known as Ci-Opsin3)(Kusakabe and Tsuda, 2007; Nakashima et al., 2003; Tsuda et al., 2003a)(**Supplemental Table 3, sheet 4**). However, expression of these and other visual cycle genes was also detected in other clusters, suggesting the presence of more than one group of molecularly distinct photoreceptor type. Indeed, photoreceptors in the *Ciona* larva have been classified, morphologically and developmentally, into three well characterized types (I, II, and III)(Horie et al., 2008b). Cluster D may correspond to only one of these three types.

### Other clusters

Cluster A closely resembled the original Cluster 5, likely comprising brain vesicle/ependymal cells (Supplemental Table 3, sheet 1), while Cluster F (Supplemental Table 3, sheet 6) appeared to be “residual” endoderm cells that did not properly cluster with clusters 1 and 6 in the original clustering.

## Discussion

By performing scRNAseq on a single batch of FACS-isolated cells at a single time-point, we have recovered transcriptional signatures of the major compartments of the *Ciona* larval brain. Although it was quite clear from our data that our FACS condition was not as stringent as we had supposed, capturing a large amount of contaminating cells from unwanted tissues such as endoderm and epidermis, scRNAseq allowed us to separate *in silico, a posteriori*, these unwanted cells from our target population. Although we may have “wasted” sequencing capacity on these unwanted cells, we now have transcriptome profiles for these cells, which may prove valuable for researchers interested in them. The separation of endoderm into 2 clusters and of epidermis into 4 clusters in our original clustering may hint at the mechanisms patterning these tissues into biologically interesting compartments.

Concerning our target tissue, the brain, the steepest and most obvious distinction we detected is that between the dorsolateral and ventral regions of the brain. The dorsolateral region contains the bulk of the differentiated neurons and photoreceptors of the brain. The ventral, proto-hypothalamo-retinal region corresponds largely to the *TH+*, dopaminergic coronet cells and perhaps related cells. These cells are largely uncharacterized, and their function in *Ciona* remains controversial. Here we have profiled their transcriptomes by both scRNAseq and bulk RNAseq, which may help guide future developmental and functional studies on this unique cell type.

The unguided PCA analysis, as currently implemented in Seurat, was able to distinguish very different cells with very different transcriptional signatures, like the coronet cells. It was not as successful to clearly identify various neuronal subtypes which undoubtedly exist in the *Ciona* brain, as deduced from their unique morphology and synaptic connectivity revealed by the connectome studies (Ryan et al., 2016, 2017, 2018). It is likely that certain subtypes are defined by expression of a few key genes, but are broadly transcriptionally similar to all other neurons in its class. It is also possible that different subtypes may be differentiated by significant variation in a larger set of low-abundance transcripts, which may not be possible to detect by scRNAseq due to technical limitations. In these cases, manual curation of scRNAseq datasets may be needed to identify hidden cell types. Furthermore, scRNAseq can still benefit from parallel analysis of more traditional bulk RNAseq analyses that rely on *a priori* selection of a large amount of cells of a single type. Perhaps the biggest caveat with our approach is its reliance on dechorionation and electroporation to label the cells for FACS. It is known that the *Ciona* larval brain is highly asymmetric (Ryan et al., 2016) and this left/right asymmetry is disrupted by dechorionation (Shimeld and Levin, 2006). For future studies, dissociation and FACS of cells from undechorionated, stably transgenic larvae (Sasakura et al., 2003) will be of utmost importance for accurate scRNAseq profiling of the *Ciona* brain.

Although the neuronal diversity within the *Ciona* brain proper remains to be elucidated, we were able to detect a small set of 12 palp neurons from the PNS, thanks in part to their distinct transcriptome and their high level of expression of the diagnostic marker *Crystallin beta/gamma*. To our knowledge, this is the first transcriptome profiling of differentiatied tunicate palp neurons performed. Accidentally extracting this transcriptome from only 12 cells in our whole dataset was only possible due to the unique advantages of the scRNAseq method and the Seurat algorithm, and suggests this strategy may be quite valuable for identifying cryptic, rare, but molecularly distinct cell types in *Ciona*. Although the palps are very important for triggering settlement and the onset of metamorphosis (Nakayama-Ishimura et al., 2009), how they do this remains virtually unknown. Perhaps the palp neuron transcriptome obtained in this study will help identify which settlement cues (e.g. tactile, chemical) they are more likely to sense and transduce.

In sum, this is the first report of using recent scRNAseq technology to profile neural tissue in *Ciona*. Our data suggest scRNAseq could be used to easily and thoroughly document transcriptome profiles from neural progenitors isolated from various stages of *Ciona* embryogenesis, and used to reconstruct the developmental patterning of the CNS. When combined with other cutting-edge techniques that have been successfully applied to *Ciona*, such as CRISPR/Cas9-mediated knockouts (Sasaki et al., 2014) and optogenetic stimulation of neural activity (Hozumi et al., 2015), scRNAseq has the potential to dramatically advance our understanding of the development and function of this minimal chordate nervous system.

## Materials and methods

### Single-cell RNAseq (scRNAseq)

Adult *Ciona robusta* (formerly *intestinalis Type A*) were collected from San Diego, CA and shipped by Marine Research and Educational Products (M-REP). Pooled eggs were dechorionated first, then fertilized with dilute sperm or 30 minutes before electroporation, as a modification of established protocols (Christiaen et al., 2009). For scRNAseq experiment, embryos were electroporated with 50 μg *Fascin>tagRFP* (see **Supplemental Sequences**) and 30 μg *Twist-related.c>eGFP* (Abitua et al., 2012) to counterselect mesenchyme cells in a final electroporation solution of 700 μl. *Twist-related.c* (KH.C5.202) was previously known as *Twist* or *Twist-like 2*. Late larvae (>20 hours post-fertilization) from 8 separate electroporations were dissociated in parallel, according to the established protocol (Wang et al., 2018). tagRFP+/eGFP^−^ cells (0.8% of the whole cell population) were isolated by FACS on a BD FACS Aria cell sorter as previously described (Wang et al., 2017; Wang et al., 2018). A 561 nm laser plus DsRed filter was used for tagRFP and a 488 nm laser plus FITC filter was used for eGFP, with nozzle size of 100 μm. Collected cells were counted and estimated to be at 63% viability using the Countess II FL automated cell counter (ThermoFisher). 10,500 viable cells were resuspended in 33.8 μl of ice-cold calcium/magnesium-free artificial seawater, 0.05% BSA and loaded into the 10X Genomics Chromium cell controller. The sequencing library was prepared using Chromium Single Cell 3’ Reagent Kit (v2 Chemistry). The library was sequenced for 100 bp paired-end reads on Illumina Hiseq 2500 system using Rapid Run mode.

### Sequencing data processing in Cell Ranger

Raw data from sequencing was processed using 10X Genomics’ Cell Ranger software according to recommended pipelines. First, the fastq files were generated using “cellranger mkfastq” with default arguments. “Read1” reads were mapped to the genome (Dehal et al., 2002; Satou et al., 2008) (http://ghost.zool.kyotou.ac.jp/datas/JoinedScaffold.zip) and counted using KyotoHoya (KH) gene models (http://ghost.zool.kyoto-u.ac.jp/datas/KH.KHGene.2013.gff3.zip) (Satou et al., 2008) by “cellranger count” with default arguments, except “--force-cells 3000”. At the same time, paired “read2” reads, which contain the cell barcode (CBC) and unique molecular identifier (UMI) sequences, were also processed by “cellranger count” to assign the counts to each cell and remove duplication artefacts introduced by library amplification.

### scRNAseq analysis in Seurat R package

We followed the procedure specified in the documentation for the Seurat scRNAseq R package (http://satijalab.org/seurat/)(Satija et al., 2015). More specific details about the exact steps can be found in the “Stage1-Clustering.Rmd” R Markdown object on our github repository (https://github.com/tunicatelab/SharmaEtAl). We filtered out low quality single cell transcriptomes by firstly, selecting genes which were expressed in at least 3 cells and selecting cells which expressed a minimum of 250 genes and a maximum of 3000 genes (**Supplemental Figure 2**). After filtering, we normalized the gene expression measurements for each cell by the total expression. This was achieved using the “LogNormalize” method. It normalizes as specified above and multiplies the value by a scale factor (10,000) and log-transforms the result. 2607 cells out of ~3000 cells passed the above quality checks and were used for downstream analysis.

### Identification of variable genes

To identify the set of genes that was most variable, we used the “FindVariableGenes” function implemented in Seurat to determine genes that were outliers on a “mean variability plot”. Average expression of each gene was calculated across all cells, and plotted against a dispersion measure for each gene. Log of variance-to-mean ratio (VMR, where average expression = mean), which is typical for counts data, was used as the dispersion measure. The genes were placed into 20 bins based on their average expression values. The dispersion measure of all genes within each bin was then z-normalized. Z-normalization allows us to compare individuals across distributions. This identified highly variable genes even when compared to genes with similar average expression. We used a z-score cutoff of 0.5 and low and high average expression cutoffs of 0.0125 and 3, respectively to identify the highly variable genes (**Supplemental Figure 3**).

### Dimensional reduction analysis

The genes identified as highly variable were then used as input for Principal Component Analysis. Before doing the PC Analysis, the data was scaled to remove the variation that could be driven by the number of detected molecules. Scaling the data could improve the downstream dimensionality reduction and clustering. PCA was then performed on the scaled data using the variable genes and the resultant PCA scores were projected onto the rest of the genes. To determine statistically significant and suitable PCs, we employed three approaches. First, we used heat maps and pairwise comparison of PCs to determine sources of variation (**Supplemental Figure 4, Supplemental Figure 5**). Second, we also used a resampling test encoded in the ‘JackStraw’ function of Seurat to identify significant PCs (**Supplemental Figure 6**). Third, we plotted the standard deviations of each of the PCs to look for a clear drop in the SDs (**Supplemental Figure 7**).

### Single cell clustering and differential gene expression

After the supervised analyses we selected principal components 1-7 for clustering and downstream analysis. To find clusters, we used the weighted Shared Nearest Neighbor (SNN) graph-based clustering method using the selected PCs. We used the “FindClusters” function with the default parameters, except for setting resolution to 0.6. We obtained 10 clusters using these. For clarity, Cluster “0” was renamed “Cluster 10” For the validation of clustering, we used the ‘ValidateClusters’ method from Seurat. We selected top 30 genes from the significant PCs and used an accuracy cutoff of 0.8 and 0.9. The minimal connectivity from the SNN was chosen to be 0.001. This, however, did not lead to merging of any of the clusters and the 10 clusters obtained initially were retained. To visualize these clusters, the cells were projected onto a 2-D plot using t-distributed Stochastic Neighbor Embedding (t-SNE) using the same 7 dimensions identified by the principal component analysis. To find differentially expressed genes among clusters, we used the “FindAllMarkers” function using the Wilcoxon Rank Sum Test (Wilcoxon, 1945) with default parameters (test.use=“wilcox”, min.pct=0.1, logfc.threshold=0.25). This selects markers as genes which are expressed in more than 10% of the cells in a cluster and average log(Fold Change) of greater than 0.25.

### Neural-specific single cell clustering and gene expression analysis

Since we were interested in neural tissue specifically, we selected a subset of the original cells for further analysis. Specifically, we selected clusters 3, 5, 8 and 9 from the first stage of clustering. We then removed all the cluster identity information from this subset and reanalyzed it using the same steps as above for identification of variable genes, dimensional reduction, clustering, and differential expression analysis with few parametric changes. More specific details about the exact steps can be found in the “Stage2-Clustering.Rmd” R Markdown object on github (https://github.com/tunicatelab/SharmaEtAl).

### Bulk RNAseq

For bulk RNAseq analysis, embryos were electroporated with 70 μg *TH>eGFP* (Moret et al., 2005), 38 μg *Fascin>tagRFP*, or 50 μg *Fascin>Unc-76::eGFP* per 700 μl electroporation solution. Late larvae were dissociated and FACS sorted as above, without counterselection directly into lysis buffer from the RNAqueous-Micro RNA extraction kit (ThermoFisher). RNA was extracted as per the kit’s manual. Two replicates of each condition (TH vs. Fascin) were prepared for cDNA synthesis as described (Wang et al., 2017), using SMART-Seq v4 Ultra Low Input RNA kit (Takara). Sequencing libraries were prepared as described (Wang et al., 2017), using Ovation Ultralow System V2 (NuGen). Libraries were pooled with other experiments and sequenced by Illumina NextSeq 500, to generate 75 bp paired-end reads. Raw sequencing fastq files were first processed by STAR 2.5.2b and mapped to the *C. robusta* genome (Dehal et al., 2002; Satou et al., 2008). The output bam files were then processed using Rsubread/featureCounts (Liao et al., 2013), with the parameter “ignoreDup=TRUE” to remove the reads duplication caused by library amplification. All the reads (after the duplication removal) which mapped to the exons of KyotoHoya (KH) gene models (Satou et al., 2008) were counted for downstream differential expression analysis. Differential expression beween TH+ and Fascin+ replicates was measured by EdgeR (Robinson et al., 2010). The exact steps can be found in the “THxFascin.Rmd” R Markdown object (https://github.com/tunicatelab/SharmaEtAl).

### In situ hybridization and reporter assays

In situ hybridization was carried out as previously described (Beh et al., 2007). *Thymosin beta-related* probe was synthesized from cloned probe template (see **Supplemental sequences**). *Fascin* reporter plasmid was constructed using a fragment upstream of the *Fascin* coding sequence, a gift kindly provided by Lionel Christiaen (see **Supplemental sequences**). Previous published drivers used in this study include: *Twist-related.c* (Abitua et al., 2012), *Tyrosine hydroxylase* (Moret et al., 2005), *Crystallin beta/gamma* (Shimeld et al., 2005), *Islet* (Wagner et al., 2014), *GAD* (Takamura et al., 2010), *Slc32a1/VGAT* (Yoshida et al., 2004), and *Slc18a/VAChT* (Yoshida et al., 2004).

### Marker gene expression search

We searched the ANISEED database (Brozovic et al., 2017) for *in situ* hybridization and other gene expression images in order to deduce the identities of the cell clusters we obtained with scRNAseq. Various images deposited on ANISEED and curated by members of the tunicate research community were embedded in Supplemental Tables 1 and 3 to highlight the expression patterns of certain genes. Links to ANISEED *in situ* pages and information on the number of images that are available for each gene on ANISEED were compiled into a single **Supplemental Table 6**, which we hope will be useful to the community for cross-referencing sequencing data with known expression patterns. Warning: the list of KyotoHoya gene model “unique names” as used in Wang et al. 2017 contains 4 entries that are often automatically, persistently converted to date format by Excel. These are “MAR5”, “MAR6”, “SEP15”, and “SEP2”. We use these names in **Supplemental Table 6**, but these names may be corrupted by Excel’s auto-format malfunction in other datasets (including our own).

## Acknowledgments

We would like to thank Lionel Christiaen for providing the *Fascin* driver and for his constant support. We are grateful to Peter Meyn for processing our samples on the 10X Genomics Chromium controller at the NYU Medical School’s Genome Technology Center. We thank Claudia Racioppi and Elijah Lowe for troubleshooting sequence analysis, and Nicholas Rouillard, Pui-Leng Ip, and Tara Rock of the NYU GenCore for advice on FACS and sequencing. S.S. is supported by a Graduate Research Assistantship in Bioinformatics from the School of Biological Sciences, Georgia Institute of Technology. This work was funded by NIH award R00 HD084814 to A.S.

